## Supplemental Figures for "Translation initiation factor eIF1.2 promotes *Toxoplasma* stage conversion by regulating levels of key differentiation factors"

[*]co-second authors

Corresponding

**Supplementary Figures**

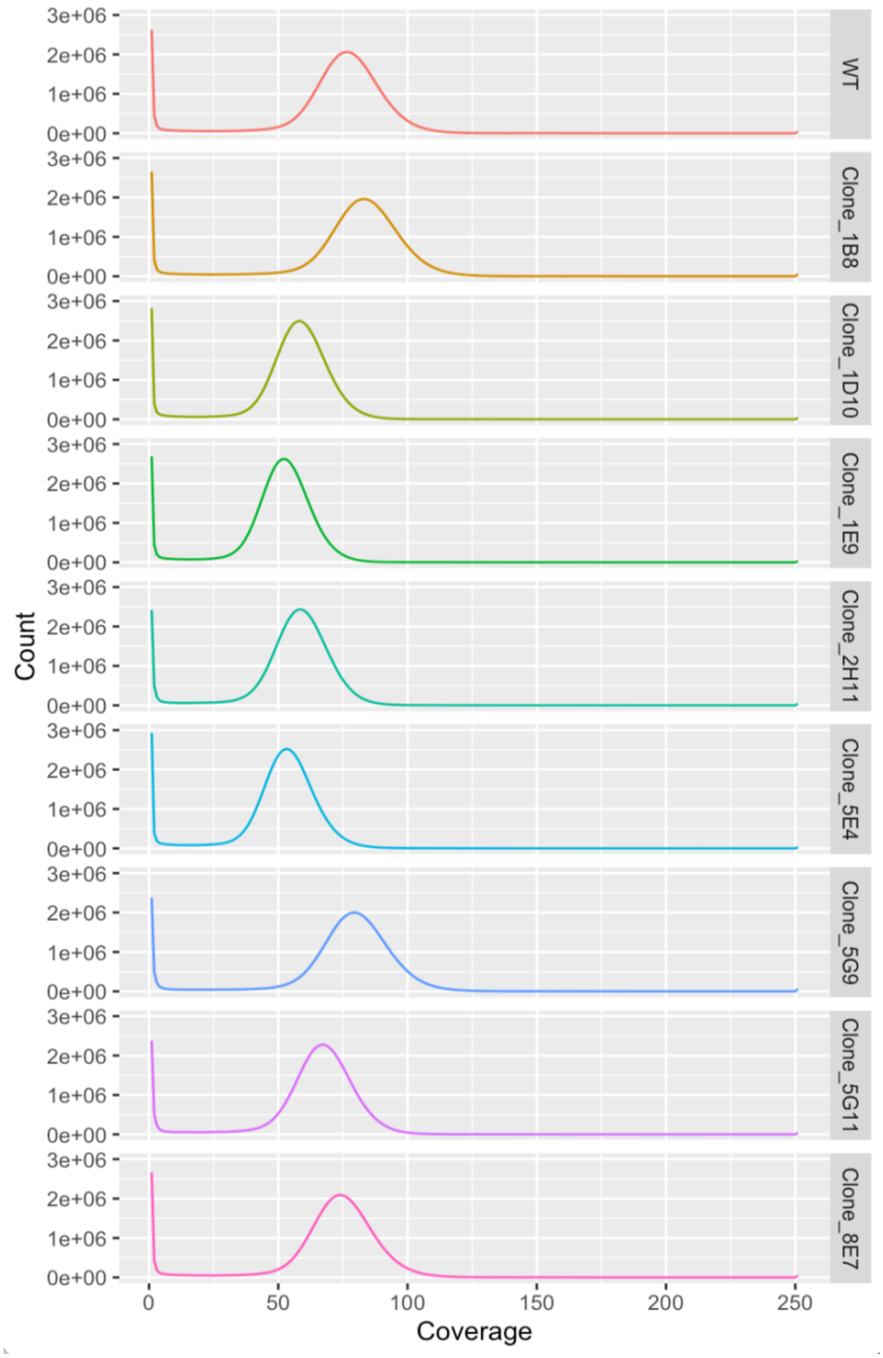


**Supplementary Fig. 1| Sequence coverage of 8 independent mutants and WT population.** These plots present the distribution of read depth across the sequenced target region. Count, the number of bases mapped the reference genome (*T. gondii* ME49). Coverage, read depth. Source data are provided as a Source Data file.

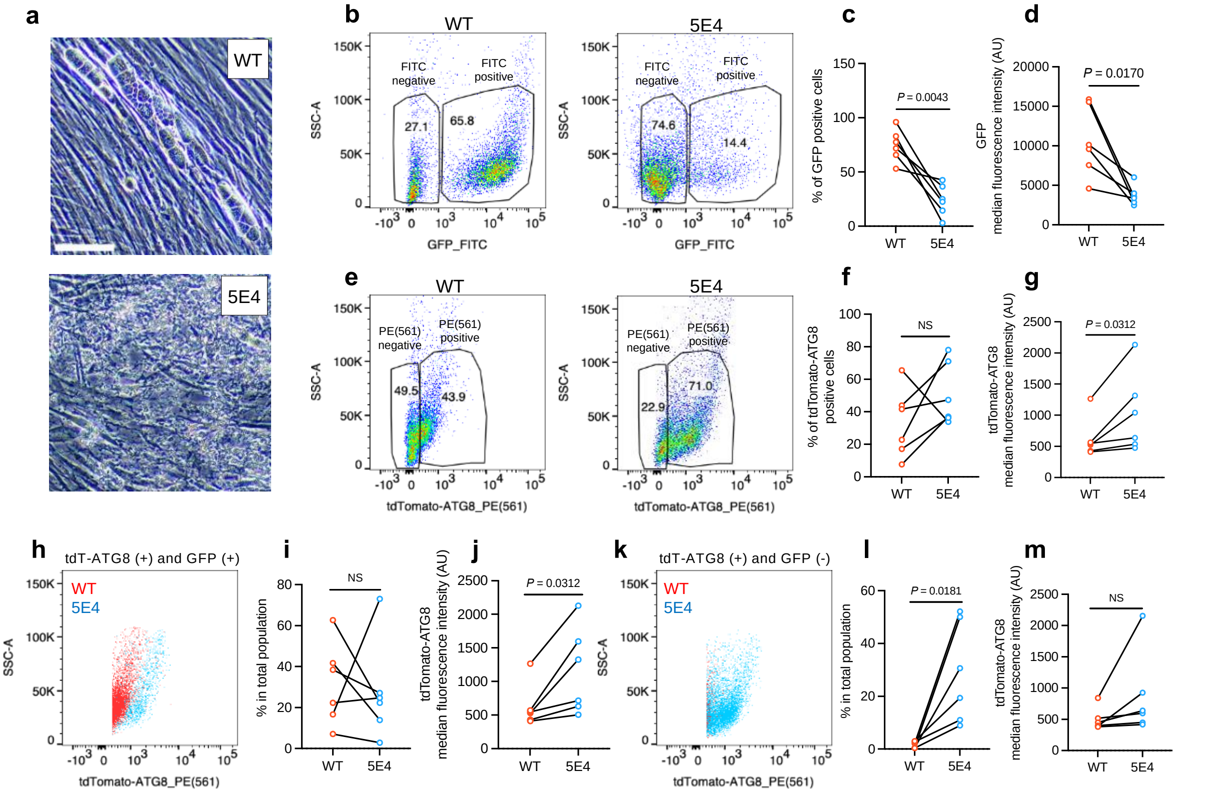


**Supplementary Fig. 2| Single mutant clone 5E4 shows a defect in differentiation.** **a**, Phase contrast microscopy images of WT (Pru ATG8) and the mutant clone 5E4 parasites after exposure to alkaline stress for 7 days. Bar, 100 μm. **b-m**, Flow analysis of parasites at 7 days post-differentiation. Experiment conducted on the same day were grouped and connected by a line. *GFP* expression is controlled by the bradyzoite-specific *LDH2* promoter. AU, arbitrary units. The numbers in the pseudocolor plots represent the percentage of the gated population relative to the total population. **b**, Representative flow cytometry pseudocolor plots of GFP fluorescence for total population. **c**, Quantification of the percentage of GFP(+) cells in total ungated population. n = 6 biological replicates; Student's two-tailed *t*-test. **d**, Quantification of GFP intensity for GFP(+) population. n = 6 biological replicates; Student's two-tailed *t*-test. **e**, Representative flow cytometry pseudocolor plots of tdTomato fluorescence for total population. **f**, Quantification of the percentage of tdTomato(+) cells in total ungated population. n = 6 biological replicates; NS, not significant ($P > 0.05$); Student's two-tailed *t*-test. **g**, Quantification of tdTomato intensity for tdTomato(+) population. n = 6 biological replicates; Wilcoxon signed rank two-tailed test. **h**, Representative flow cytometry pseudocolor plots of tdTomato fluorescence for tdTomato(+) and GFP(+) population. **i**, Quantification of the percentage of tdTomato(+) and GFP(+) cells in total ungated population. n = 6 biological replicates; NS, not significant (*P* > 0.05); Student's two-tailed *t*-test. **j**, Quantification of tdTomato intensity for tdTomato(+) and GFP(+) population. n = 6 biological replicates; Wilcoxon signed rank two-tailed test. **k**, Representative flow cytometry pseudocolor plots of tdTomato fluorescence for tdTomato(+) and GFP(-) population. **l**, Quantification of the percentage of tdTomato(+) and GFP(-) cells in total ungated population. n = 6 biological replicates; Student's two-tailed *t*-test. **m**, Quantification of tdTomato intensity for tdTomato(+) and GFP(-) population. n = 6 biological replicates; NS, not significant (*P* > 0.05); Student's two-tailed *t*-test. Source data are provided as a Source Data file.

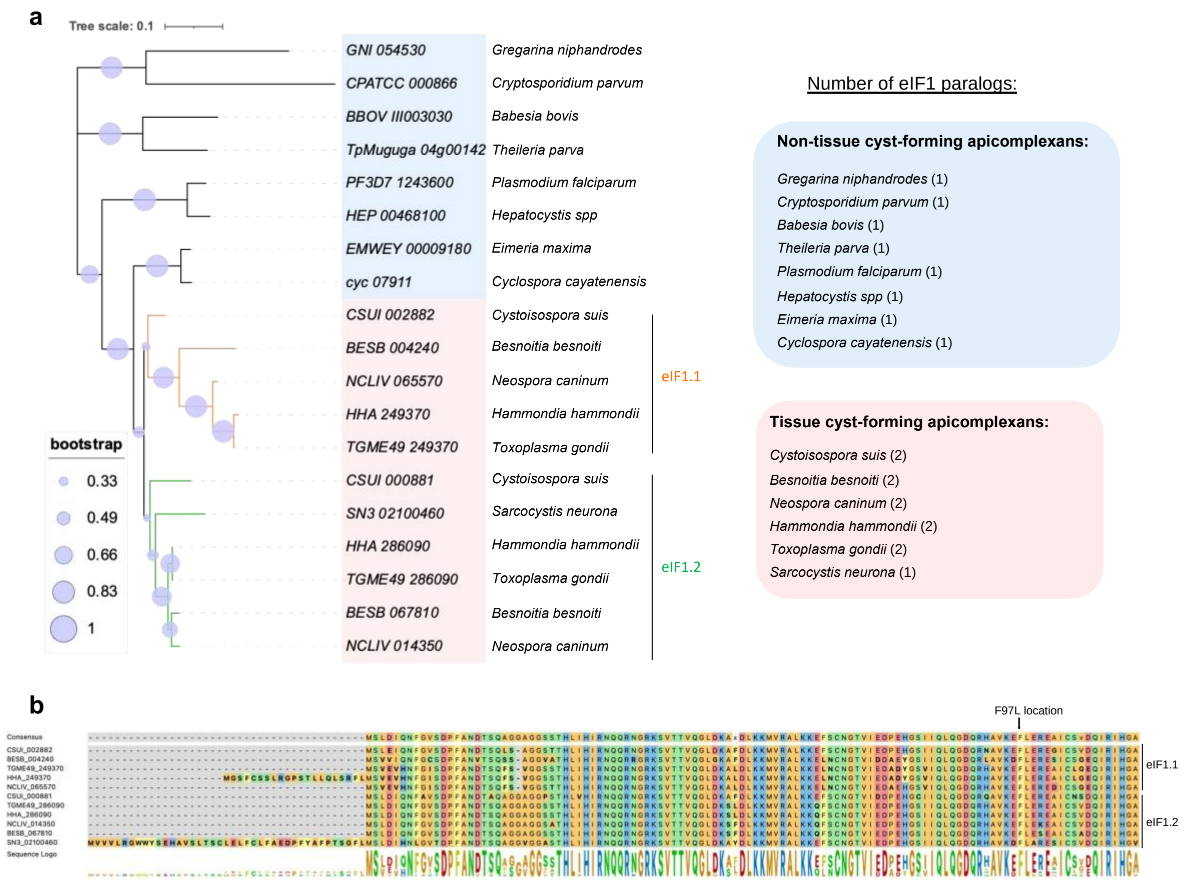


**Supplementary Fig. 3|eIF1 paralogs in apicomplexan parasites. a**, Neighbor-joining phylogenetic tree of eIF1 proteins for apicomplexan parasites. Green branches represent proteins that share a greater similarity to eIF1.2 (TGME49_286090), while orange branches correspond to proteins closely related to eIF1.1 (TGME49_249370). Clades: blue (non-tissue cyst-forming coccidia), pink (tissue cyst-forming coccidia). Scale bar represents the number of amino acid substitutions per site along a branch. Bootstrap values were computed based on 1000 replications. **b**, Alignment of these eIF1.1 and eIF1.2 proteins from **a** using CLustalW. The location of F97L mutation in eIF1.2 is indicated.

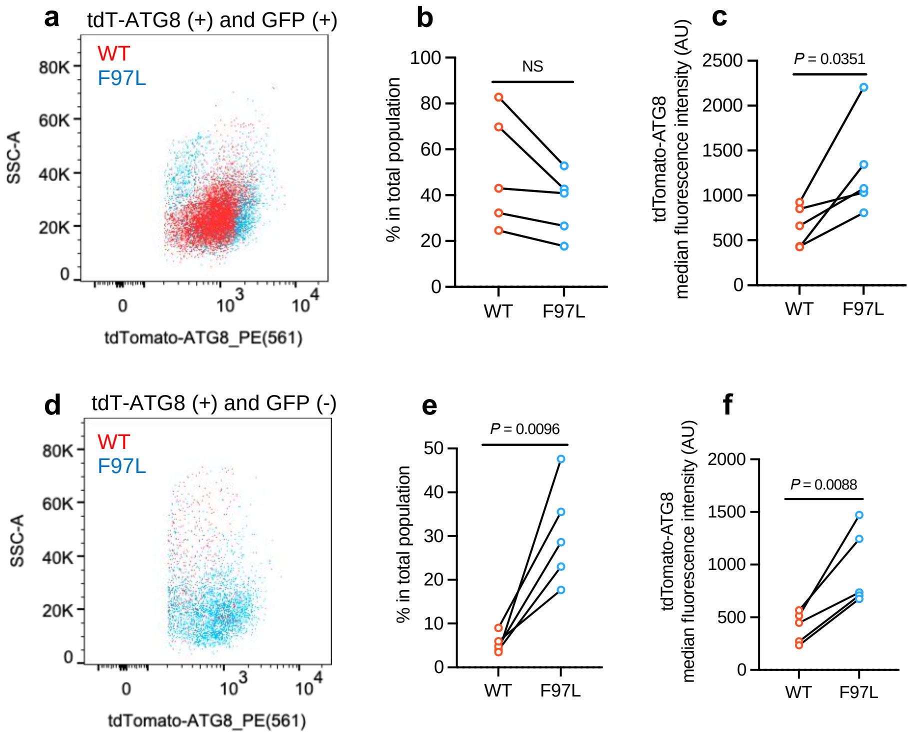


**Supplementary Fig. 4|The differentiation defect of eIF1.2 F97L mutant is not correlated with increased tdT-ATG8 intensity, as related to Fig. 2. a-f,** Parasites exposed for 7 days to alkaline stress. Experiment conducted on the same day were grouped and connected by a line. AU, arbitrary units. **a**, Representative flow cytometry pseudocolor plots of tdTomato fluorescence for tdTomato(+) and GFP(+) population. **b**, Quantification of the percentage of tdTomato(+) and GFP(+) cells in total ungated population. n = 5 biological replicates. NS, not significant ($P > 0.05$); Student's two-tailed *t*-test. **c**, Quantification of tdTomato intensity for tdTomato(+) and GFP(+) population. n = 5 biological replicates; Student's two-tailed *t*-test. **d**, Representative flow cytometry pseudocolor plots of tdTomato fluorescence for tdTomato(+) and GFP(-) population. **e**, Quantification of the percentage of tdTomato(+) and GFP(-) cells in total ungated population. n = 5 biological replicates; Student's two-tailed *t*-test. **f**, Quantification of tdTomato intensity for tdTomato(+) and GFP(-) population. n = 5 biological replicates; Student's two-tailed *t*-test. Source data are provided as a Source Data file.

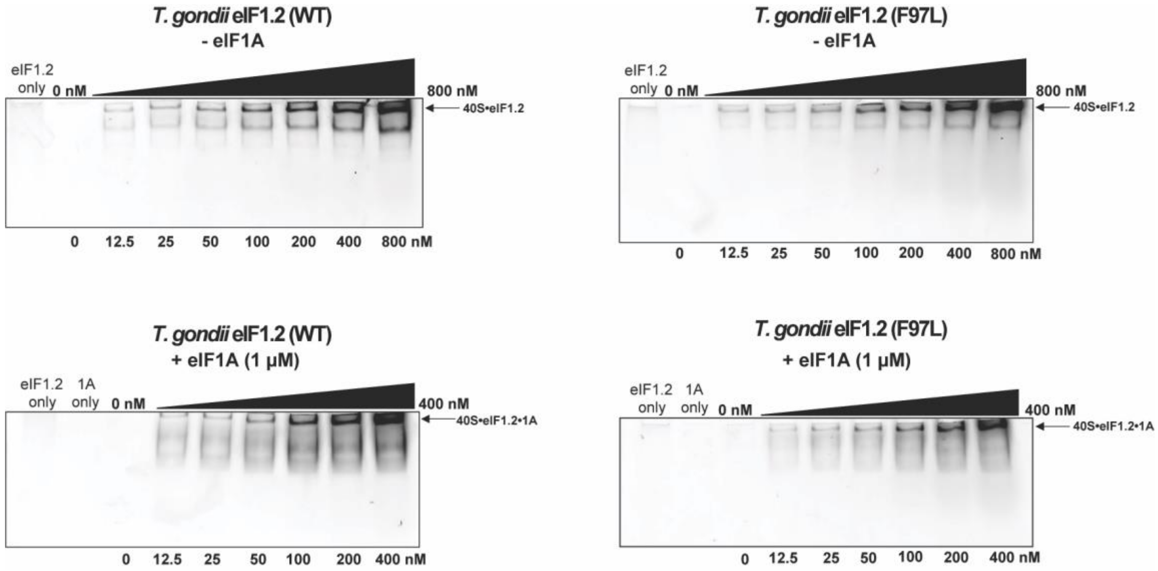


**Supplementary Fig. 5|Gel shift analysis of *T. gondii* eIF1.2 WT or F97L binding to yeast 40S, as related to Fig. 3.** Representative gels illustrate yeast 40S (100 nM) binding to varying concentrations of *T. gondii* eIF1.2 (12.5 to 800 nM in the absence of yeast eIF1A; 12.5 to 400 nM in the presence of yeast eIF1A (1 μM)). The lane labeled "eIF1.2 only" contains 100 nM eIF1.2. Source data are provided in Supplementary Information file.

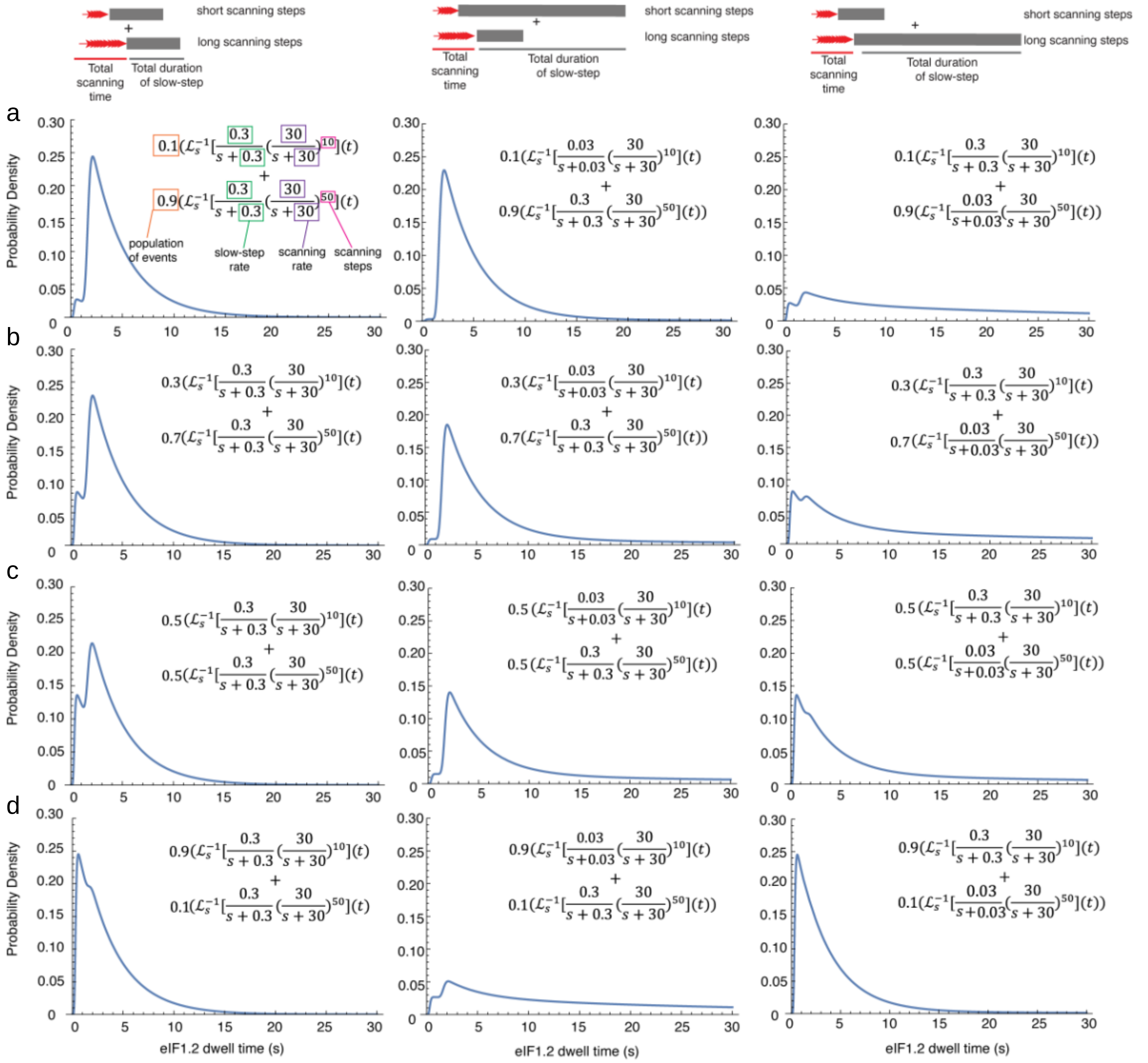


**Supplementary Fig. 6| Scanning Simulation indicates eIF1.2 F97L does not decelerate a slow step.** Simulated eIF1.2 dwell-time probability density functions reflect two independent, parallel processes: 10 and 50 scanning steps, modeled as irreversible processes. The 10-step process reflects PIC recognition of an upstream start site, 10 nucleotides from the mRNA 5' end. The 50-step process represents scanning bypass of the upstream site, with subsequent recognition of a downstream start site positioned 50 nucleotides from the mRNA 5' end. Each process includes an additional slow step, modeling eIF1.2 ejection. Probability density functions were generated by an inverse Laplace transform approach[1]. The relative prevalence of each pathway was systematically varied by coefficients multiplying the individual density functions (increasing relative prevalence of the 10-step pathway from top to bottom). To understand whether the increased eIF1.2 dwell time population for eIF1.2(F97L) was due to a deceleration of a slow step in the scanning process (e.g., eIF1.2 ejection), we reduced the slow-step rate (initially 0.3 $s^{-1}$) by 10-fold in either the short (10) or long (50) dwell time populations. **a**. 10% short+90% long. **b**. 30% short+70% long. **c**. 50% short and 50% long. **d**. 90% short and 10% long. Reducing the slow-step rate introduced a dominant, quasi-exponential component to the distribution. This behavior was not observed in our single-molecule scanning experiments. In the single-molecule scanning experiments, the positions, and shapes of the short- and long-time components do not change, but their relative contributions to the overall distribution changed with the F97L substitution.

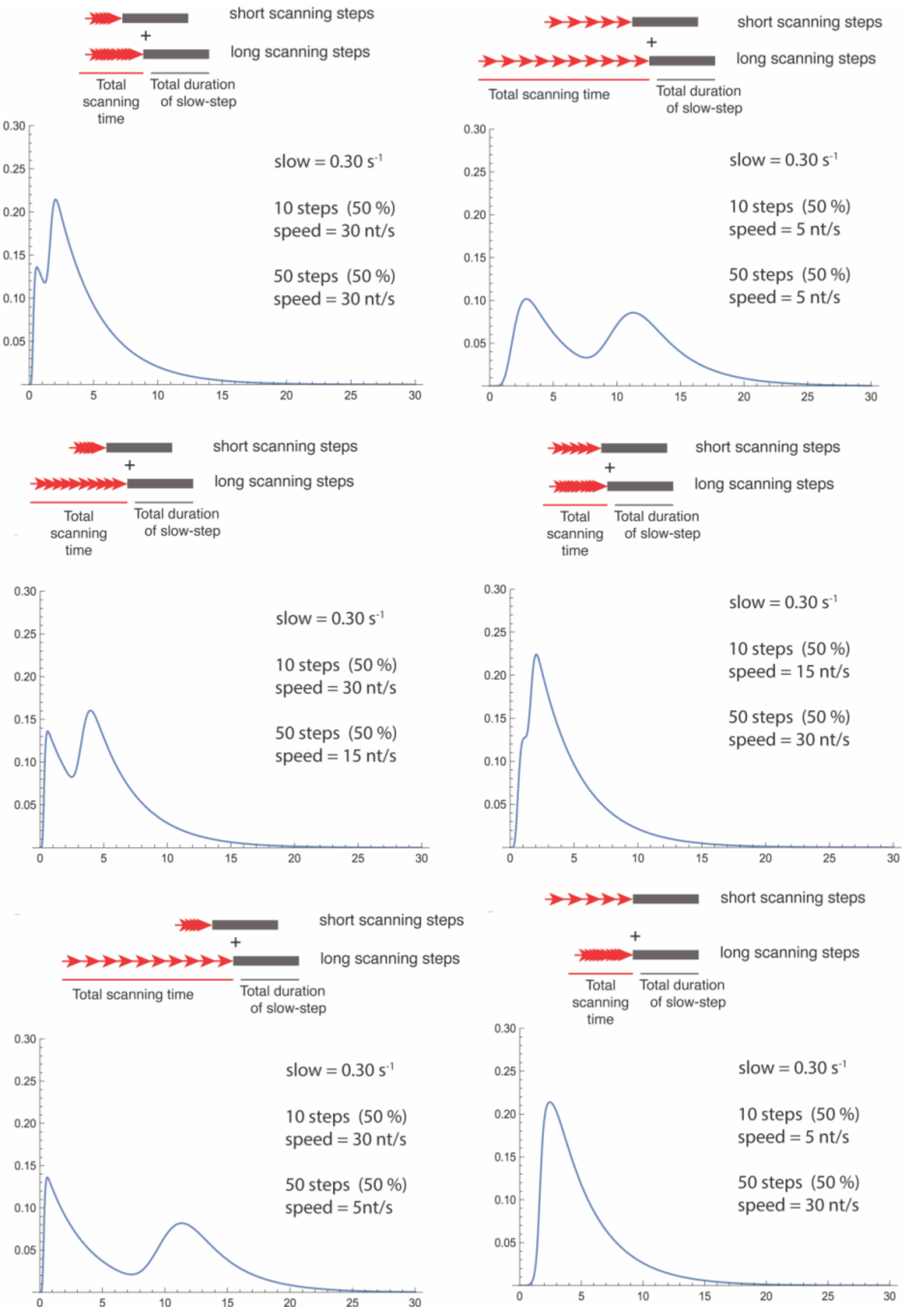


**Supplementary Fig. 7|Scanning simulation suggests that eIF1.2 F97L does not impede the scanning rate.** Simulated eIF1.2 dwell-time probability density functions reflect two independent parallel processes: 10 and 50 scanning steps, modeled as irreversible processes. The 10-step process reflects PIC recognition of an upstream start site, 10 nucleotides from the mRNA 5' end. The 50-step process represents scanning bypass of the upstream site, with subsequent recognition of a downstream start site positioned 50 nucleotides from the mRNA 5' end. Each process includes an additional slow step, modeling eIF1.2 ejection. Probability density functions were generated by an inverse Laplace transform approach[1]. The relative scanning rates for each pathway were systematically varied, while the slow eIF1.2 ejection rate remained constant; the relative prevalence of each pathway was equal in each simulation. The simulated results showed that slowing scanning rate for short and long components shift the positions of their respective distributions to longer times, which was not observed in our single-molecule scanning experiments. In our single-molecule scanning experiments, the positions, and shapes of the short- and long-time components do not change, but their relative contributions to the overall distribution changed with the F97L substitution.

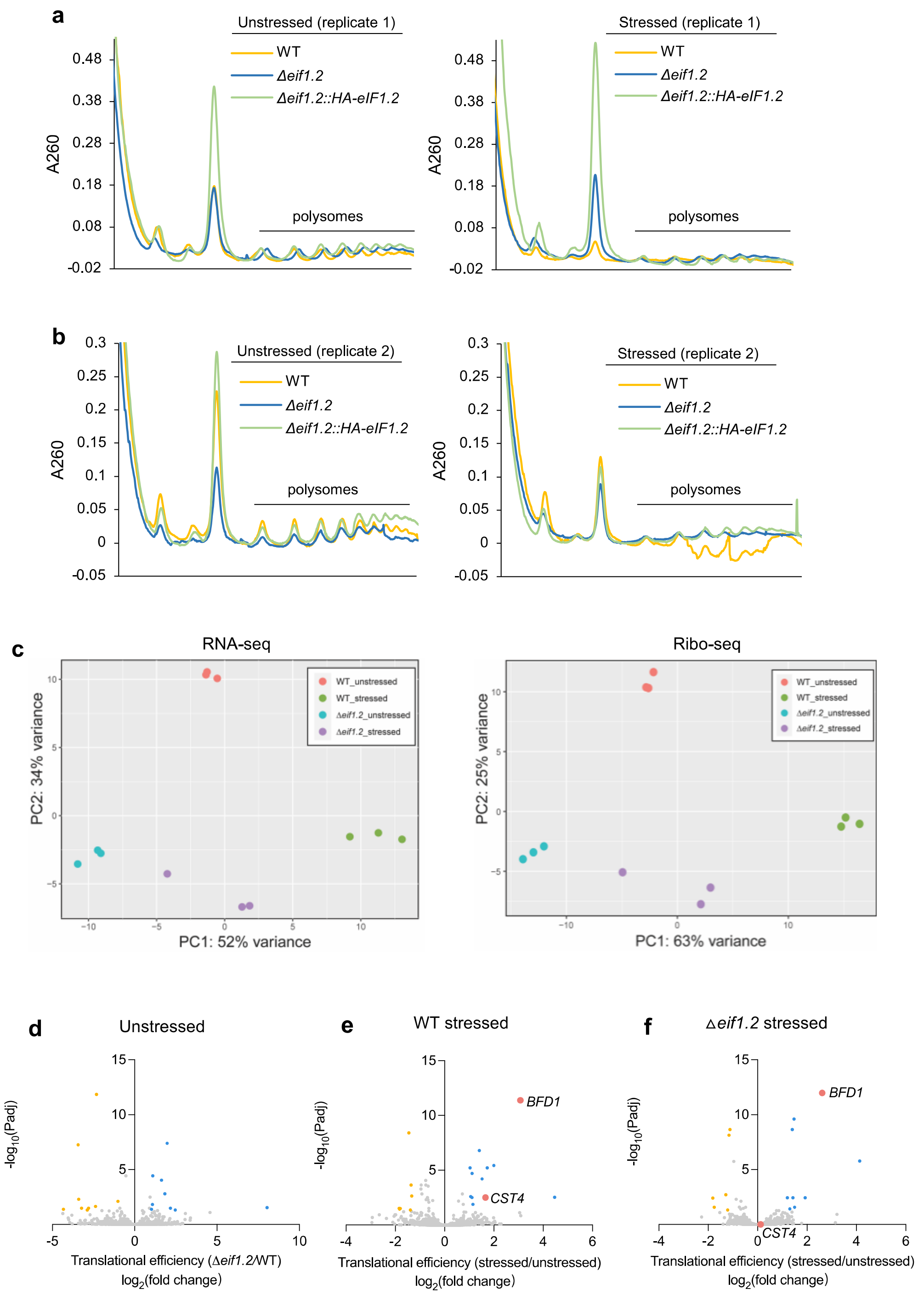


**Supplementary Fig. 8|Polysome profiling, analyses of RNA sequencing and ribosome profiling, as related to Fig. 5. a**,**b**, Polysome profiling experiments. n = 2 biological replicates. Data were adjusted during plotting to align the 80S peak location of all samples with that of unstressed WT samples, and the lowest A260 absorbance values around the 80S peak for all samples were set to 0. **c**, Principal component analysis for RNA sequencing (RNA-seq) and ribosome profiling (Ribo-seq) data. Stressed, 1 day of alkaline stress. WT, ME49Δ*ku80*. n = 3 biological repeats. **d**, Differential analysis of translational efficiency in Δ*eif1.*2 parasites versus WT parasites under unstressed conditions. **e**, Differential analysis of translational efficiency in WT parasites under stressed versus unstressed conditions. **f**, Differential analysis of translational efficiency in Δ*eif1.*2 parasites under stressed versus unstressed conditions. Source data are provided as a Source Data file.

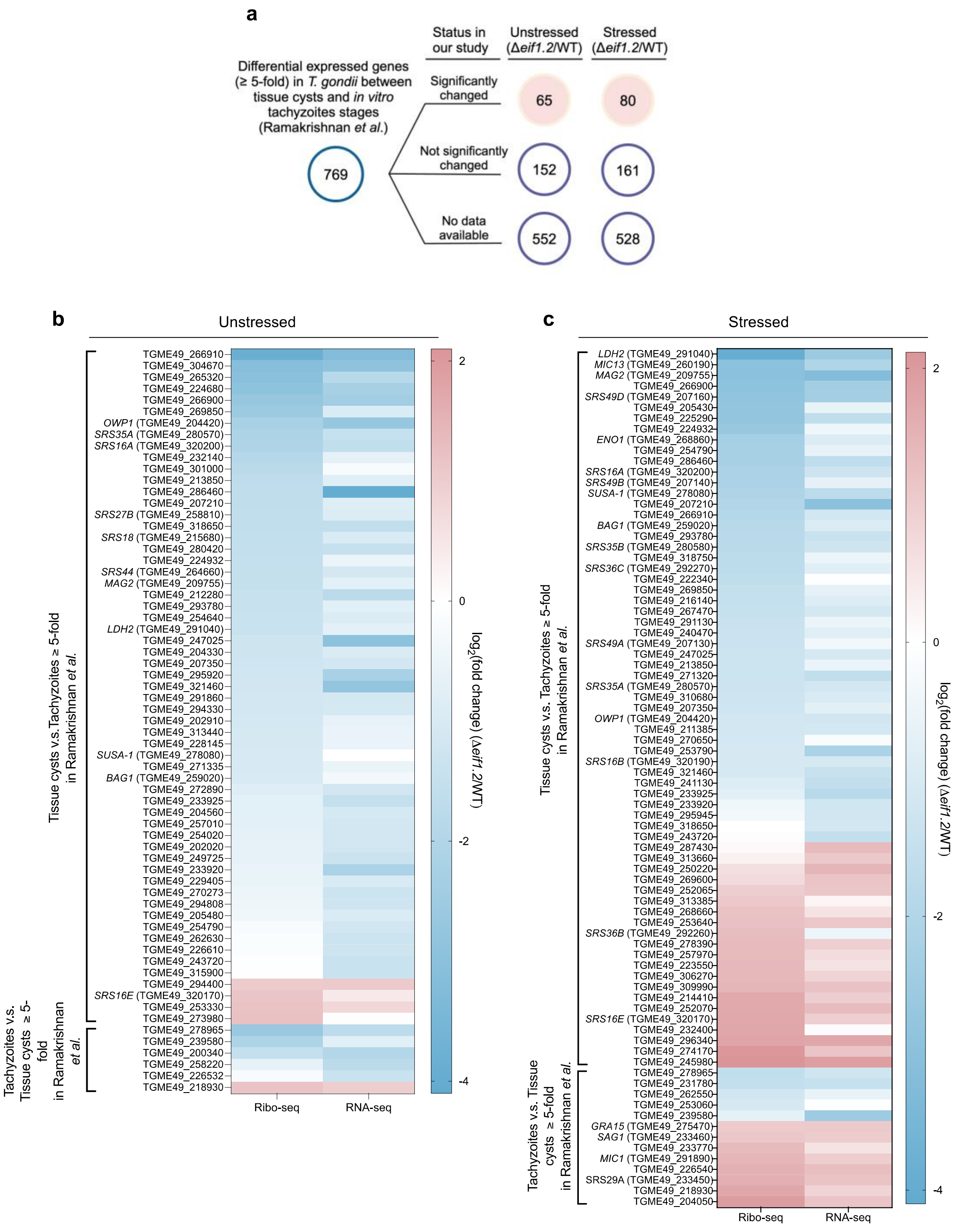


**Supplementary Fig. 9| a-c**, Comparative analysis of our RNA-seq and Ribo-seq results with a previously published RNA-seq dataset for tissue cysts and *in vitro* tachyzoites[2]. Gene listed as significantly changed in our study have a fold change greater than 2 or less than 0.5 in either RNA-seq, Ribo-seq, or both, with $P_{adj} < 0.05$, and minimum of 5 reads. **b**,**c,** Heatmaps depict the fold change of genes significantly altered between unstressed Δ*eif1.2* and WT parasites (**b**, 65 genes), and between stressed Δ*eif1.2* and WT parasites (**c**, 80 genes) in our study ($P_{adj}$ < 0.05). Source data are provided as a Source Data file.

### Supplementary References

1. Floyd, D. L., Harrison, S. C. & Van Oijen, A. M. Analysis of Kinetic Intermediates in Single-Particle Dwell-Time Distributions. *Biophys. J.* **99**, 360–366 (2010).
2. Ramakrishnan, C. *et al.* An experimental genetically attenuated live vaccine to prevent transmission of Toxoplasma gondii by cats. *Sci. Rep.* **9**, 1474 (2019).

**Source Data**

### Supplementary Fig. 5

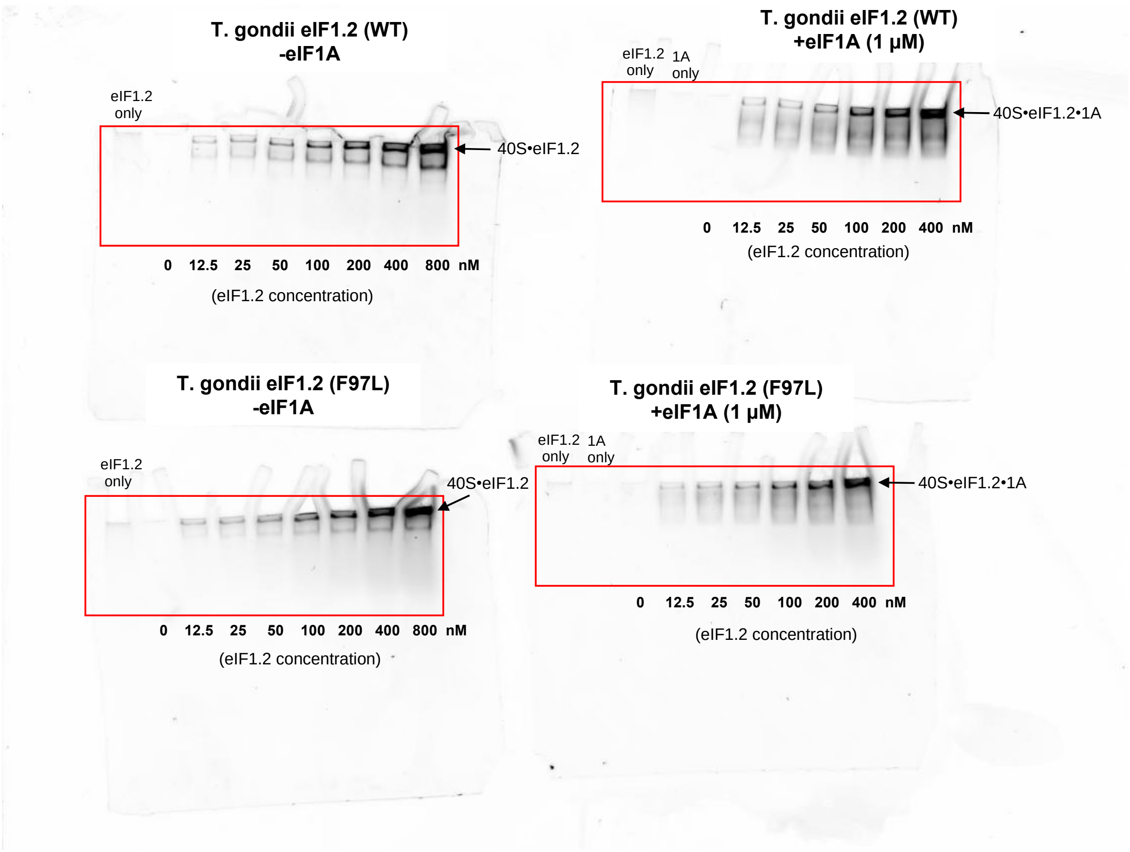
